# Predicting drug targets by homology modelling of *Pseudomonas aeruginosa* proteins of unknown function

**DOI:** 10.1101/2021.06.15.448521

**Authors:** Nikolina Babic, Filip Kovacic

## Abstract

Efficacies of antibiotics to treat bacterial infections rapidly decline due to antibiotic resistance. This stimulated the development of novel antibiotics, but most attempts failed. As a response, the idea of mining uncharacterised genes of pathogens to identify potential targets for entirely new classes of antibiotics raised. Without knowing the biochemical function of a protein it is difficult to validate its potential for drug targeting; therefore progress in the functional characterisation of bacterial proteins of an unknown function must be accelerated. Here we present a paradigm for comprehensively predicting biochemical functions of a large set of proteins encoded by hypothetical genes in human pathogens, to identify candidate drug targets. A high-throughput approach based on homology modelling with ten templates per target protein was applied on the set of 2103 *P. aeruginosa* proteins encoded by hypothetical genes. Obtained >21000 homology modelling results and available biological and biochemical information about several thousand templates was scrutinised to predict the function of reliably modelled proteins of unknown function. This approach resulted in assigning, one or often multiple, putative functions to hundreds of enzymes, ligand-binding proteins and transporters. New biochemical functions were predicted for 41 proteins whose essential or virulence-related roles in *P. aeruginosa* were already experimentally demonstrated. Eleven of them were shortlisted as promising drug targets which participate in essential pathways (maintaining genome and cell wall integrity), virulence-related processes (adhesion, cell motility, host recognition) or antibiotic resistance, which are general drug targets. These proteins are conserved among other WHO priority pathogens but not in humans, therefore they represent high-potential targets for pre-clinical studies. These and many more biochemical functions assigned to uncharacterised proteins of *P. aeruginosa*, available as PaPUF database may guide the design of experimental screening of inhibitors which is a crucial step toward validation of the most potential targets for the development of novel drugs against *P. aeruginosa* and other high-priority pathogens.

## Introduction

### Last minutes for radical changes in discovering novel treatments for bacterial infections

Even after more than a half-century development of antibiotics, bacterial infections remain a significant cause of morbidity and mortality worldwide because of unsuccessful antibiotic treatments. The rise in multidrug-resistance bacterial infections has been further accelerated by the overuse of antibiotics in present pandemic conditions. [1] It is evident that without new solutions against multidrug-resistant bacteria, so-called “superbugs”, routine medical procedures (cancer chemotherapy, organ transplantation, diabetes treatment) become very high risk and the Millennium Development Goals’ achievement is endangered. [2]

Attempts of the World Health Organisation (WHO) to guide and promote research and development (R&D) of new antibiotics failed. [2, 3] The hurdles faced by the current system for R&D of antibiotics are tremendous, what is the reason that only a few solutions (32 antibiotics and ten biological treatments) against the WHO priority pathogens are currently in the R&D pipeline (clinical phases 1 to 3). [2–4] Generally, the most attractive targets for new classes of antibiotics are proteins involved in previously untargeted essential biochemical pathways conserved among pathogens and absent in humans. [1, 5] Targeted screening of such enzymes identified reductase IspH involved in isoprenoid biosynthesis [6], reductase FabI involved in fatty acid biosynthesis [7] and peptidase LspA involved in posttranslational processing of lipoprotein [8] which respective inhibition by C23.28–TPP (pre-clinical study) [6], isoniazid (approved drug) [7] and myxovirescin (clinical phase 3) [8] showed strong antibacterial activity.

New antibiotics represent only one solution with uncertain efficacy because of the fast resistance evolution observed after each new antibiotic was introduced. [9] Another promising strategy is led by the idea of counteracting the non-essential functions mediated by bacterial virulence or antibiotic resistance factors, thereby blocking the host-pathogen interactions, the progress of infection or antibiotic expelling without killing it or inhibiting its growth. Such antivirulence drugs suggested as a promising supplement with enhancing effects on classical antibiotics are thought to have weak selective pressure minimizing the risk of resistance emergence. [10] Among the main antivirulence approaches with drugs in clinical development are inhibition of quorum-sensing systems, biofilm formation, and toxin production or function. [7, 10]

Obviously, several proteins targeted by novel potential antibiotics or antivirulence drugs were identified in the last decade though; many more novel essential proteins or important virulence factors with unknown function in bacteria still need to be characterised as it is difficult to set up an inhibitor screening for proteins of unknown function (PUFs). [1] Therefore, understanding the function of these proteins is a prerequisite for the validation of targets for the development of antibacterial drugs with novel mechanisms of action.

### P. aeruginosa is a notorious and important human pathogen

*P. aeruginosa*, a critical priority pathogen according to WHO, is a serious threat in hospitals where multidrug-resistant strains cause severe acute infections, often with deadly outcomes. [3] Life-threatening chronic *P. aeruginosa* infections were frequently observed in immunocompromised persons, e.g. cystic fibrosis, cancer patients. [11] Concern about *P. aeruginosa* infections is owed to the high resistance towards currently the best available antibiotics - carbapenems and the third generation of cephalosporins. [3] To cope with antibiotics, *P. aeruginosa* uses various intrinsic and acquired resistance mechanisms including expression of different classes of β-lactamases, increased outer membrane impermeability by losing outer membrane channel proteins, overexpression of antibiotic efflux pumps, synthesis of antibiotic modifying enzymes and mutations in primary targets of quinolone antibiotics. [12] Moreover, the formation of antibiotic-resistant biofilms [12], exceptional adaptability to infect various human tissues owing to multiple signalling pathways [13] and cell surface adhesins [14, 15] and fast progress of infections as a result of the production of a vast number of potent extracellular toxins [16] challenge the treatment of *P. aeruginosa* infections. All these features qualify *P. aeruginosa* as a model pathogen bacterium for the R&D of novel antibiotics.

Here we aimed to assign the biochemical functions to hypothetical proteins from a human pathogen *P. aeruginosa*. Using homology modelling-based high-throughput approach, novel putative functions were reliably assigned to many proteins for which functions could not be previously predicted from the sequence. Several proteins with new putative biochemical functions could be linked to important infection-related processes what suggests that these proteins are promising targets for the development of drugs against *P. aeruginosa*.

## Material and Methods

Gene and protein sequences were retrieved from the *Pseudomonas* genome database [17] (PGD, www.pseudomonas.com) and Uniprot database [18] (www.uniprot.org), respectively. PGD annotations were used to find proteins having human homologs, proteins having homologs in other pathogens and virulence factors. The list of virulence factors was fulfilled by using the Virulence factor database (www.mgc.ac.cn/VFs) [19] and Victors virulence factors knowledge database (www.phidias.us/victors) [20]. Comparative BLAST was performed using NCBI’s BLAST suite (https://blast.ncbi.nlm.nih.gov/Blast.cgi). Homology modelling was performed using the batch processing feature of the Phyre2 web server [21, 22] Selected models were several times energy minimized with the GROMOS96 43B1 force field as implemented in Swiss-PdbViewer [23]). The stereochemical parameters of the homology models were evaluated using the PROCHECK v.3.5 program [24] DALI database [25] (http://ekhidna2.biocenter.helsinki.fi/dali/oldstyle.html) was used to identify structures similar to homology models. The protein structures were analyzed, aligned and visualized using PyMol software (http://www.pymol.org). A Diamond BLASTp tool [26] from PGD was used to search for groups of orthologs with >70% sequence identity and >95% alignment coverage, among different *P. aeruginosa* strains isolated from humans with developed disease as indicated in PGD and from other pathogenic *Pseudomonas* species, namely *P. alcaligenes*, *P. mendocina,* and *P. otitidis*. NCBI BLAST was used to search homologs with >35% sequence identity and >75% alignment coverage in 13 WHO priority pathogens.

## Results

### *P. aeruginosa* genes of unknown function are potential antibiotic targets

Development of efficient drugs against multiresistant bacteria relies on the identification of entirely new bacterial targets whose inhibition will hamper bacterial growth (antibiotic) or prevent pathogen from being virulent (antivirulence drugs). Such targets are likely to be found among genes of unknown function (GUF), so-called genomic ‘dark matter’. [1] Interestingly, even 20 years after the *P. aeruginosa* PA01 genome was sequenced, nearly 40% of genes are still GUFs and this is a current state in the majority of sequenced bacterial genomes. [27, 28]

To identify GUFs in *P. aeruginosa* PAO1, we were searching functional class IV hypothetical genes in manually curated *Pseudomonas* Genome Database (PGD). [17] Results revealed 1470 hypothetical genes, which have no homologs in other species and 633 conserved hypothetical genes, which have homologs without known function in other bacteria (Fig. 1A and Table S1). These 2103 genes of *P. aeruginosa* considered as GUFs were analysed by integrating comparative sequence analysis and structural homology-based functional prediction.

**Fig. 1.**
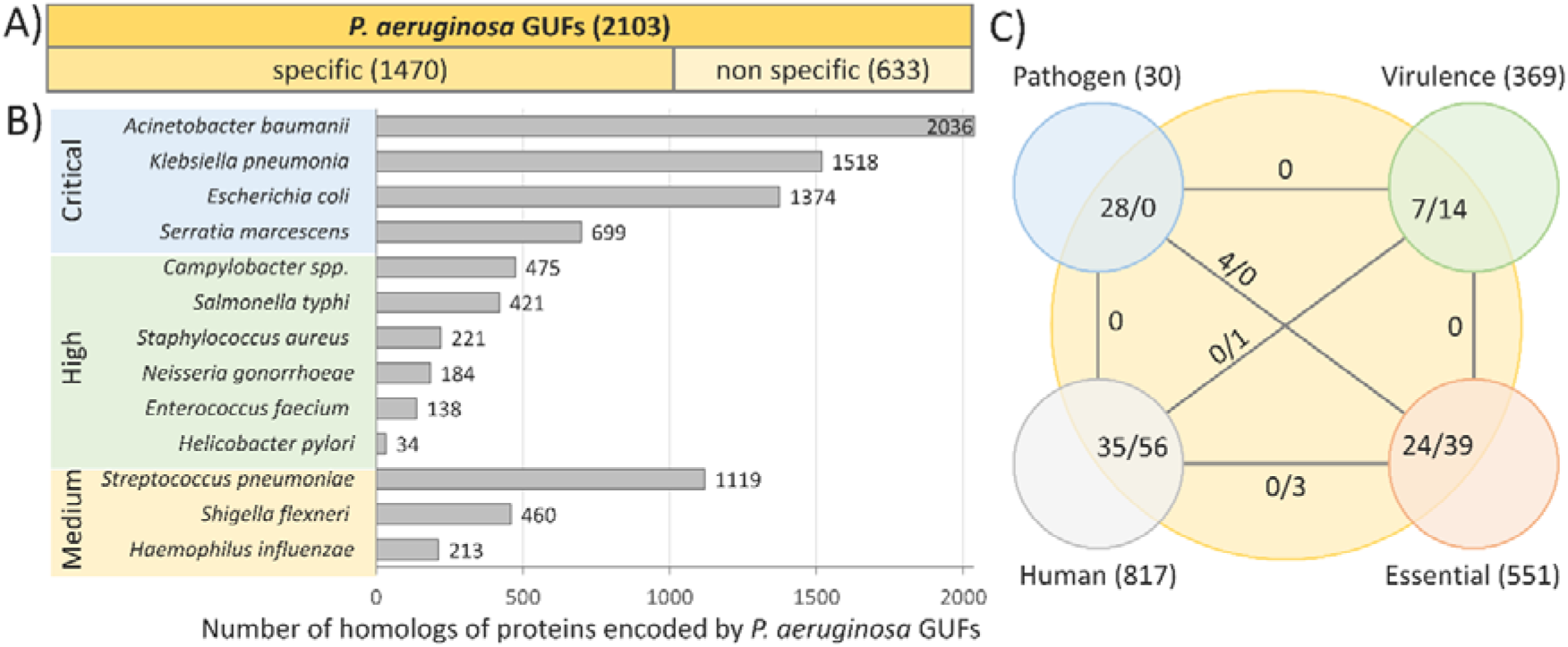
A large portion of genomes of *P. aeruginosa* PA01 and WHO priority pathogens consist of GUFs. **A)** Majority of *P. aeruginosa* GUFs were found only in *Pseudomonas* species (specific) while some of them are conserved in other organisms (non-specific). The analysis was performed using the PGD version **20.2** download on 5.11.2020. **B)** Conservation of *P. aeruginosa* genes of unknown function in WHO priority pathogens. *P. aeruginosa* PUFs were used as a query for BLAST search on NCBI to find protein homologs (>35% sequence identity, >75% sequence coverage) in following thirteen WHO priority pathogens. Taxonomic identifiers of organisms are listed in Table S2. **C)** Pathogen-specific, essential, virulence factors and genes with human homologs among GUFs. Numbers in brackets indicate the number of genes in each group. A number of *P. aeruginosa* GUFs in overlapping sections is indicated as a number of specific genes/ number of non-specific genes. Numbers on lines indicate mutual GUFs among two linked groups.

Bacterial antibiotic targets absent in humans are particularly interesting, as their inhibition should not exert side effects on host physiology. [1, 7] Using PGD [17] annotation of *P. aeruginosa* proteins having human homologs we have shown that the majority of PUFs (2012) do not have human homolog (Fig. 1C).

The essential genes, genes conserved only in pathogenic species and genes with demonstrated virulence function represent high priority target-groups for R&D of antibiotics because they may participate in key cellular or virulence processes whose inhibition, might be lethal or virulence-attenuating. [29] We found that 63 GUFs of *P. aeruginosa* belong to essential genes (Fig. 1C and Table S1), identified as those genes whose experimental mutation is leading to cell death. [30] Furthermore, overlapping *P. aeruginosa* GUFs with *P. aeruginosa* genes which have homologs only in pathogenic microorganisms (pathogen-associated genes) [31] revealed 28 hits. Four of them (PA0442, PA0977, PA1369, PA2139) are particularly promising targets for broad range antibiotics as they were also annotated as generally essential genes (Fig. 1B and Table S1).

Furthermore, we have analysed which GUFs might play a role in the pathogenic processes by searching virulence-associated GUFs listed in virulence factor annotation of PGD. [17] In this list are genes whose roles in virulence was experimentally proven, by studying respective mutant strains, or was inferred from the sequence similarity with virulence factors from other pathogens. Even 13 identified GUFs were previously linked to virulence-related process by studying their functions during infection of model host organisms (e.g. *Caenorhabditis elegans*, *Drosophila melanogaster* or *Rattus norvegicus*) and additional five were related to common virulence traits (flagella-mediated motility, pyoverdin-mediated iron uptake, type VI secretion of toxins) (Table S3). [32–36] Among these virulence-associated GUFs, eleven are conserved in other organisms (Table S3), suggesting that their analysis might reveal novel insights in general virulence processes in bacteria.

### Assigning the putative biochemical functions to *P. aeruginosa* proteins of unknown function

To better understand the role of GUFs during infections with *P. aeruginosa,* it is necessary to know the biochemical functions of proteins encoded by GUFs. We have used homology modelling of a 3D structure as a tool for predicting biochemical functions of PUFs. This approach relies on the fundamental principle that two homologous proteins with similar sequences have very similar structures. As protein structure is 3 – 10 times more conserved than the sequence, [37] structural modelling can reliably detect homology between proteins with low sequence similarity (<20%). [21] Knowing that structure determines protein function, one can assume that the functions of modelled PUF and the template protein with a known structure are likely to be the same. This approach used the advantage of, in most cases, precisely assigned biochemical and even biological, functions to proteins with known structure. A further advantage is that generated structural models of target PUFs are useful for validation of functionally relevant residues, e.g. active site or ligand binding residues. Additionally, several distinct functional domains in single target PUF may be identified using different domains with solved structures and these domains can be combined into a unique multi-domain structure. Conclusively, homology modelling of PUF and its comparison with the template may reliably reveal the putative function of PUF.

Using protein sequences of PUFs obtained from the Uniprot database, we have performed homology modelling of 2103 PUFs using in batch homology modelling feature of the Phyre2 server. [21] Retrieving ten models per PUF we have compiled a set of data about 21020 (PA2462 with a size of 573.2 kDa was too large for homology modelling) homology modelled templates (Table S4). These results were refined to select reliable models using an automated pipeline (Fig. 2). A Phyre confidence score, which indicates a degree of homology between target proteins and the template, was used as the main criterion for validation of structural prediction. According to this parameter for most PUFs (75%), fairly reliable homology models (>80% conf.) were obtained (Fig. S1A), although their sequence identities were rather low since almost 65 % of modelled proteins had sequence identities below 20% compared with corresponding templates (Fig. S2B). Interestingly, 89 PUFs have very low confidences (<20%) (Table S1), which is an indication that these PUFs do not have even remote structural homolog. Hence, these proteins might be either intrinsically disordered or have a novel fold and therefore are particularly interesting from the structural biology point of view. Interestingly, we identified 65 PUFs with known structure as their sequence identities with template were >96%. Among them were 18 proteins with experimentally demonstrated functions (Table S5); therefore, they were excluded from our further analysis and the information is now uploaded to PGD to improve its annotation.

**Figure 2:**
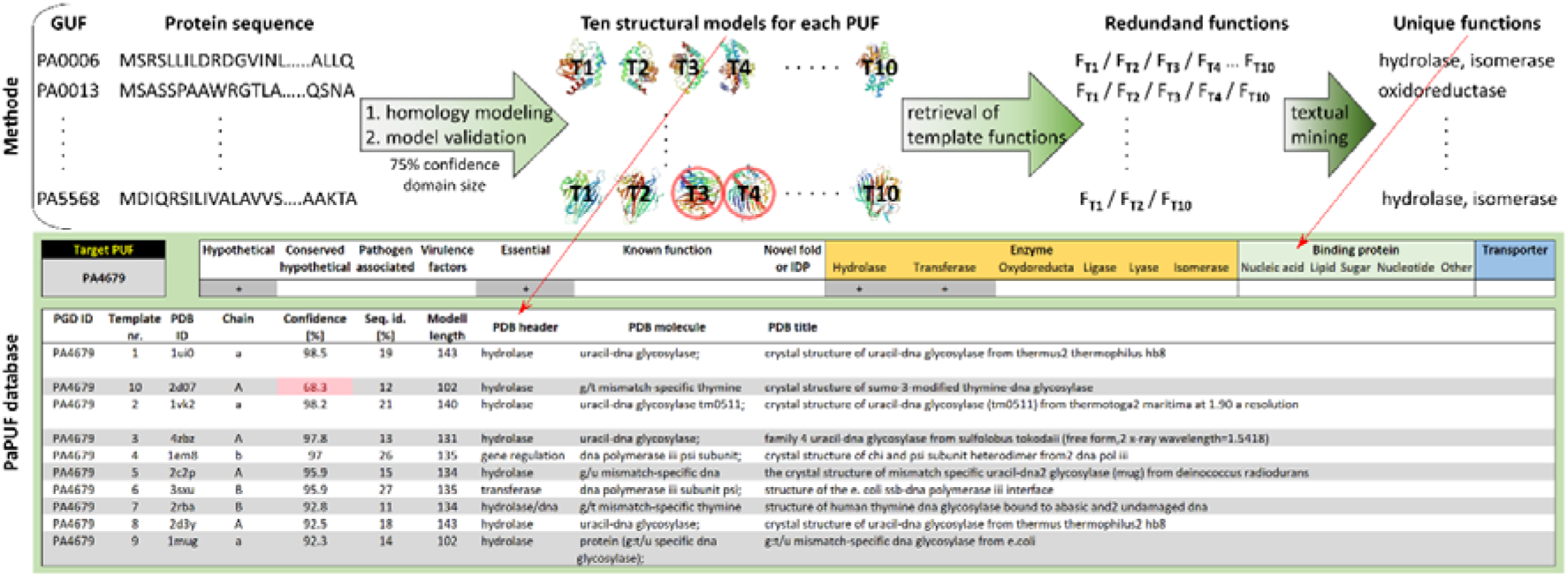
Pipeline for prediction of protein function based on homology modelling with multiple templates. Protein sequences of all 2103 *P. aeruginosa* GUFs were retrieved from the UniProt database and submitted to the Phyre2 server to model their structures. Ten structural models for each PUF were validated according to the confidence of homology modelling (>75%) and the size of the modelled domain (>100 amino acids for transporters and enzymes; >50 amino acids for binding proteins) to remove unreliable models. Information about the functions of reliable templates was retrieved from their PDB header and PDB molecule fields and text mining with keywords listed in table S6 was performed to assign biochemical functions to templates automatically. Redundant functions were removed to compile a list of unique functions of templates which were assigned as putative functions of PUFs. Results are freely available as downloadable PaPUF database (www.iet.uni-duesseldorf.de/arbeitsgruppen/bacterial-enzymology) in which putative functions of *P. aeruginosa* PUFs and respective ten templates can be retrieved by entering a gene name into the “Target PUF” field (upper left corner).

To predict the biochemical functions of PUFs with the potential to be novel drug targets, we have focused on enzymes, ligand binding proteins and transporters, which are carriers of functions indispensable for bacterial cell cycle, infections and antibiotic resistance, e.g. primary metabolism, toxin-mediated host cell lysis, cell surface adhesion and the antibiotic expelling. [15, 38, 39] Consequently, their inhibition might lead to death or reduce the virulence of *P. aeruginosa*. To increase the reliability of functional assignment, we have eliminated homology models smaller than 100 residues for enzymes and transporters and smaller than 50 residues for ligand-binding domains. This is in agreement with a thermodynamic folding study which suggests that the optimal size of protein domains is 100 residues. [40] The cut-off for rejection of binding domains was set to 50 residues as generally a helix-turn-helix zinc finger DNA-binding domains, [41] a five-stranded beta-barrel oligonucleotide and oligosaccharide-binding domains [42] and lipid-binding domains, [43] of virulence-related proteins, are usually not smaller than 50 residues. This comprehensive and accurately refined data set consisting of, mostly, several reliable protein templates per PUF was a starting point for functional prediction using the information about biochemical functions of templates. General text mining algorithm based on the set of keywords (Table S6) which precisely describe here studied protein functions resulted in the prediction of enzyme, binding or transport functions for 59% (1228) PUFs (Fig 3A). Interestingly, among them were 304 PUFs with two and 59 with three of here analysed functions (Fig 3B).

**Figure 3:**
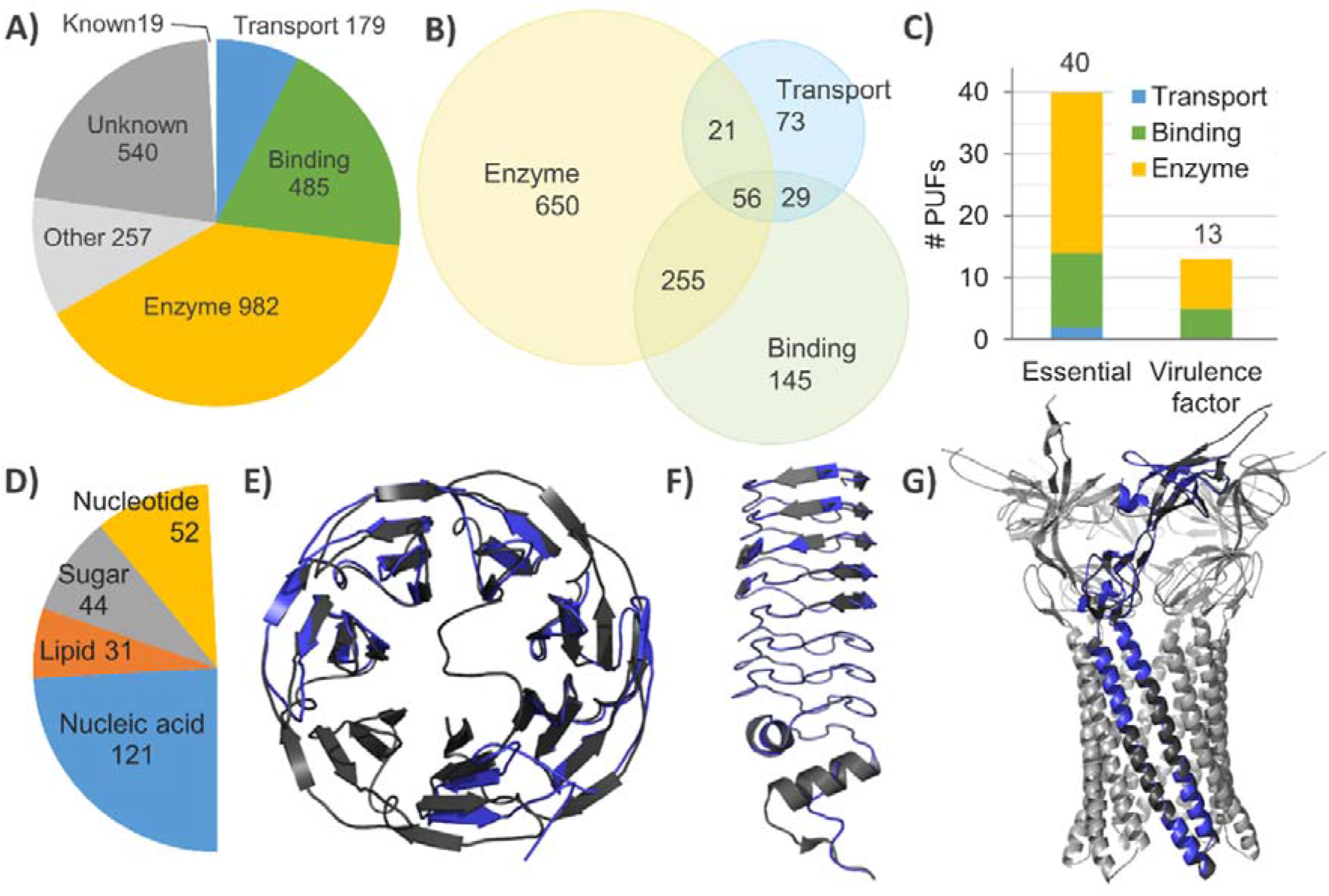
Predicted biochemical functions for *P. aeruginosa* proteins of unknown function. **A)** Number of putative enzymes, ligand-binding proteins, transporters, proteins with other functions than previous three and proteins with known functions. **B)** Venn diagram of PUFs with an enzyme, ligand binding a transport functions. **C)** Distribution of essential and virulence factors among PUFs with the predicted enzyme, ligand binding or transport functions. Gene identifiers of PUFs with here predicted functions (Figs. 3A – C) are provided in table S1. **D)** Number of predicted ligand-binding proteins. **E)** Superimposition of a homology model of putative lectin PA5033 with the *A. aegerita* lectin (PDB ID 4TQJ, chain A, 88.4% conf., 19% seq. id., 1.11 Å rmsd_cα_, 92% coverage, 1.3 % of disall. res.). **F)** Superimposition of a homology model of putative DNA gyrase binding-protein PA1981 with the *Mycobacterium tuberculosis* MfpA protein (PDB ID 2BM5, chain A, 99% conf., 22% seq. id., 1.45 Å rmsd_cα_, 80% coverage). **G)** Superimposition of a homology model of putative antibiotic efflux pump protein PA3304 with the *E. coli* MacA (PDB ID 3FPP, chain A, 99% conf., 23% seq. id., 0.56 Å rmsd_cα_, 81% coverage, 1.9 % disall. res.). PA3304 is superimposed to one of six MacA protomers, which form a barrel-like functional protein. A homology model is indicated blue and corresponding template grey. The protein structures were visualised using the Pymol software.

Unfortunately, the functions of four proteins that belong in *Pseudomonas*-specific and essential protein groups (Fig. 1C) were not predicted. Analysis of enzymes, transporters and binding proteins with newly predicted functions revealed that even 41 GUFs might be considered as potential drug targets as they were found in essential or virulence-related groups (Fig. 3C). Interestingly comparative sequence analysis reveals that these proteins have 400 homologs (>35% seq. id.) in priority pathogens (Table S2). Among them are only four proteins with human homologs (Table S2). Thus, here presented assignment of putative functions illustrates the global significance for R&D of potential bacteria-specific drug targets.

### Virulence-related functions of putative ligand-binding proteins and transporters

Ligand binding domains were predicted in 484 proteins, which showed a large overlap (310 proteins) with putative enzymes (Fig. 3B). Among them, the proteins predicted to bind nucleic acids were the most abundant (122) followed by nucleotide-binding (52), sugar-binding (44) and lipid-binding (33) proteins (Fig. 3D). An interesting example of virulence-related binding protein is putative lectin PA5033 (Fig. 3E), which was reliably modelled (88% conf.) with the structure of lectin from a mushroom *Agrocybe aegerita*. [44] Reliability of homology modelling is furthermore supported by only a small portion (1.3 %) of residues in disallowed (disall. res.) regions of Ramachandran plot as revealed by the stereochemical validation of energy minimised PA5033 homology model with PROCHECK (Table S7). [24] Hence, the most (~80%) high-quality X-ray structures have <5 % residues in disallowed regions. [45] This protein with experimentally determined extracellular localisation may be involved in bacterial adhesion to host cells or in biofilm formation as described for the other two (LecA and lecB) extracellular lectins of *P. aeruginosa*, which were suggested as targets of biofilm-specific antivirulence drugs. [46] The further interesting candidate is putative DNA gyrase binding protein PA1981 (Fig. 3F) which was reliably (99% conf., 0.6 % disall. res.) modelled using the *Mycobacterium tuberculosis* MfpA protein. [47] Proteins from this family bind to DNA gyrase, an antibiotic target with the essential function for the homeostatic control of chromosomal DNA supercoiling, thereby protecting it against antibiotic fluoroquinolone. Therefore, PA1981 is suggested as a protein implicated in resistance against fluoroquinolone.

An example of antibiotic-resistance protein is putative membrane-bound transporter PA3304 that was reliably modelled with *E. coli* MacA adaptor protein of efflux pump involved in antibiotic expelling (Fig. 3G, Table S7). [48] How many of 105 putative membrane transporters, among 179 in this study predicted transporters (Fig. 3A), play role for antibiotic resistance, tolerance to xenobiotics or secretion of virulence factors remains to be elucidated.

### Putative enzymatic activities of hypothetical proteins link them to important cellular processes

Putative enzymes accompany the largest group (982) of PUFs. Their functional predictions might help to identify potential novel drug targets involved in fundamental metabolic pathways of *P. aeruginosa*. The classification of enzymes using text mining revealed hydrolases as the most abundant class, followed by transferases and oxidoreductases (Fig. 4A). Thirty-five of PUFs with putative enzyme functions are particularly interesting drug targets as they belong to a group of essential proteins or virulence factors (Fig. 4B). Ten of them have two and one even three predicted enzyme activities what indicates profound substrate promiscuity among potential drug targets. An interesting candidate with essential function is PA4679, a putative uracil-DNA glycosylase (UDG) whose function is predicted based on the modelling with *Thermus thermophiles* UGD (Fig. 4C). The functional prediction of UDG activity of PA4579 is strengthened by the finding that the structure of the homology model is similar to several other enzymes with UDG activity (root mean square deviation of Cα atoms, rmsd_cα_ < 1.7 Å) and active site residues of *Tt*UGD interacting with the uracil molecule are conserved among them (Figs. 4C and S3, Table S8). Its essentiality is in agreement with the function of these enzymes for DNA base excision repair, a process essential for maintaining the stability of chromosomal DNA. [49]

**Figure 4:**
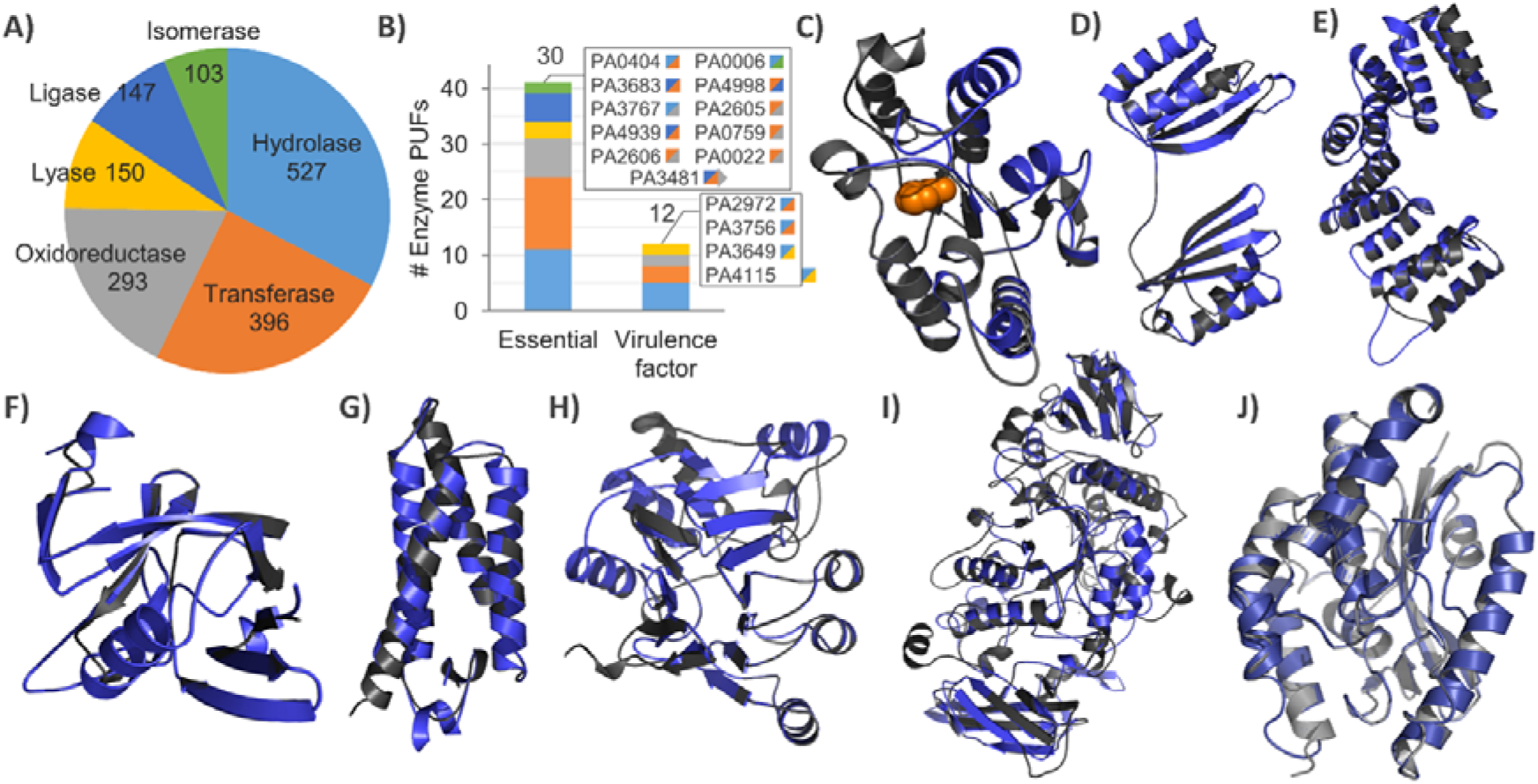
Putative enzymatic activities of *P. aeruginosa* proteins of unknown function. **A)** Distribution of putative enzymes to six EC groups. **B)** Distribution of essential and virulence factors among GUFs with putative enzyme functions. Gene identifiers of PUFs with here predicted functions (Figs. 4A and B) are provided in table S1. **C)** Superimposition of a homology model of putative uracil-DNA glycosylase PA4679 with the *T. thermophiles* UGD (PDB ID 1UI0, chain A, 98.5% conf., 19% seq. id., rmsd_cα_ = 0.62 Å, 61% coverage, 0.9 % disall. res.). The uracil molecule bound to the active site groove of *Tt*UGD is shown as an orange space model. **D)** Superimposition of a homology model of putative phosphoserine phosphatase PA1009 with the *M. avium* SerB (PDB ID 3P96, chain A, 99% conf., 12% seq. id., rmsd_cα_ = 0.50 Å, 94% coverage, 1.2 % disall. res.). **E)** Superimposition of a homology model of putative Type IV pili biogenesis proteins PA5441 with the *N. meningitides* PilW (PDB ID 2VQ2, chain A, 93% conf., 17% seq. id., rmsd_cα_ = 0.29 Å, 38% coverage, 0.5 % disall. res.). Residues 1 – 130 of the template are hidden for the purpose of representation. **F)** Superimposition of a homology model of putative L,D-transpeptidases PA3756 with the *M. tuberculosis* LdtMt2 (PDB ID 3VYN, chain B, 100% conf., 18% seq. id., rmsd_cα_ = 0.65 Å, 83% coverage, 0.8 % disall. res.). **G)** Superimposition of a homology model of putative flagellar chaperon PA1095 with the *A. aeolicus* FliS (PDB ID 1ORJ, chain A, 100% conf., 28% seq. id., rmsd_cα_ = 0.46 Å, 94% coverage, 0% disall. res.). **H)** Superimposition of a homology model of putative phosphorylcholine esterase PA2984 with *S. pneumoniae* Pce (PDB ID 1WRA, chain A, 100% conf., 16% seq. id., rmsd_cα_ = 0.40 Å, 32% coverage, 3% disall. res.). **I)** Superimposition of a homology model of putative glycoside hydrolase PA2151 with *S. pneumoniae* SpuA (PDB ID 2YA0, chain A, 100% conf., 17% seq. id., rmsd_cα_ = 0.54 Å, 94% coverage, 1.9 % disall. res.). **J)** Superimposition of a homology model of putative patatin-like phospholipase PA1640 with *P. aeruginosa* outer membrane phospholipase PlpD (PDB ID 5FYA, chain A, 100% conf., 33% seq. id., rmsd_cα_ = 0.36 Å, 72% coverage, 1.4% disall. res.). A homology model is indicated blue and corresponding template grey. The protein structures were visualised using the Pymol software (http://www.pymol.org).

We have predicted biochemical functions of four PUFs for which the role in virulence was previously experimentally determined (Table S3). Putative hydrolase PA1009 was reliably modelled using a template of phosphoserine phosphatase SerB from *Mycobacterium avium* which is involved in a pyrimidine salvage pathway important for genome integrity [50] (Fig. 4D, Table S7). C-terminal part of putative cell motility protein PA5441 was reliably modelled with Type IV pili biogenesis proteins PilW from the *N. meningitides* which exerts an important role for oligomerisation and functionality of outer membrane-localised pore of pili [51] (Fig. 4E). Putative transferase PA3756 is modelled with 100% confidence and good stereochemical parameters (0.8 % of disall. res.) using *M. tuberculosis* L,D-transpeptidase (LdtMT2), which is a common target of β-lactam antibiotics [52] (Fig. 4F and Table S7). The putative function of PA1095 as a FliS-like chaperon, assigned from the homology with *Aquifex aeolicus* FliS (Fig. 4G) [53] involved in the assembly of the bacterial flagellum. Virulence function prediction for PA5441, PA1009, PA3756 and PA1095 is in agreement to their virulence role experimentally determined in genetic screening of mutants in *Rattus norvegicus* virulence assay [54] or motility assay [36].

Furthermore, we have predicted the functions of several PUFs based on the homology to known virulence factors from other pathogens. Although these proteins were not previously related to the virulence in *P. aeruginosa*, their putative biochemical functions suggest that they might be novel virulence factors of *P. aeruginosa*. Among them are three enzymes, PA2984, PA2151 and PA1640, with putative hydrolytic activities. The C-terminal part (residues 494 – 736) of PA2984 was reliably modelled (Fig. 4H, Table S7) with phosphorylcholine esterase Pce from *Streptococcus pneumoniae,* whose virulence function is related to the adhesion to host cells. [55] The N-terminal domain of PA2984 could not be modelled with any known structure; however, the analysis with Phyre2 predicted 12 putative transmembrane helices in this part (Fig. S2), thus suggesting that this putative hydrolase is a two-domain protein in which catalytic domain is anchored to the membrane by an N-terminal domain. Interestingly, the *S. pneumoniae* template protein has also a second domain which through binding of surface choline exposes the esterase domain to the cell surface. The structure of putative hydrolase PA2151 was reliably modeled (1.9 % of disall. res.) (Fig. 4I, Table S7) with glycogen-degrading virulence factor, SpuA from *Streptococcus pneumoniae*, which is involved in host-pathogen interactions. [56] Putative extracellular hydrolyse PA1640 was modelled (Fig. 4J) with 100% confidence (1.4 % of disall. res.) using the structure of type V secreted phospholipase PlpD which is described as putative virulence factor of *P. aeruginosa* [57, 58]. PlpD belongs together with *P. aeruginosa* type III secretory toxin ExoU [59] to the patatin-like family of phospholipases. Patatin domain-containing proteins are very potent toxins due to their strong lytic activity which they exert on host cells through, among others, hydrolysis of host membrane phospholipids [59].

The functions of the above-mentioned enzymes could not be predicted from the sequence alignment as they share less than 20% sequence identities with the template proteins (except PA1095 which share 28% seg. id.). However, high confidence of homology modelling and good stereochemical parameters of the homology models demonstrate that homology modelling may be used even to predict the biological process in which protein is involved.

### Conservation of promising drug targets in other pathogens

Protein conservation in multiple pathogens is an important criterium used for the prioritization of targets for the development of drugs. The broad-range drugs have an advantage over the pathogen-specific drugs because of higher profits, easier development due to uncomplicated recruitment of patients for phase III trials and no requirement of specific diagnostic tools for pathogen detection. [2] Therefore, we have searched orthologs of promising drug targets in clinical isolates of *P. aeruginosa*, in other pathogenic *Pseudomonas* species (*P. mendocina*, *P. otitidi* and *P. alcaligenes*) and in pathogens from WHO priority list (Table 1). Results revealed that each of eleven promising targets with newly predicted ligand binding (Fig. 3) or enzyme (Fig 4) functions are conserved (>70% sequence identity, >95% alignment coverage) in most clinical isolates of *P. aeruginosa* (Table 1). Furthermore, ten of eleven promising targets are conserved (>70% sequence identity, >95% alignment coverage) in at least two important pathogens from *Pseudomonas* genus. Finally, most of these eleven proteins are broad-range targets as all of them have homologs (the nine of eleven proteins were conserved in four or more pathogens) in other WHO priority pathogens (Table 1). Interestingly, even seven potential drug targets have homologs in Gram-negative and Gram-positive bacteria (Table 1) indicating that a drug binding these targets could have very broad spectrum. Hence, these proteins whose function was previously unknown do not have human homologs (Table 1) what make them even more attractive for development of drugs against the infections with *Pseudomonas* and other important bacterial pathogens.

**Table 1:** Promising drug targets of *P. aeruginosa* among proteins of unknown function.

| Gene | Putative biochemical function | Putative virulence function | Phyre 2 confidence [%] | Conservation |  |  |  |
| --- | --- | --- | --- | --- | --- | --- | --- |
|  |  |  |  | WHO priority pathogens<sup>§</sup> | pathogenic *Pseudomonas*<sup>€</sup> | clinical *P. aeruginosa*<sup>£</sup> | humans |
| PA5033 | lectin | cell adhesion | 88 | *Ab, Kp* | *Po, Pm* | 2519 | no |
| PA1981 | DNA gyrase binding-protein | antibiotic resistance | 99 | *Ab, Sp, Ec, Sm, Sa, Kp, St* | *Pm* | 261 | no |
| PA3304 | efflux pump adapter protein | antibiotic resistance | 99 | *Ab, Kp, Ec, Sf, Sm* | *Po, Pm, Pa* | 2523 | no |
| PA4679 | uracil-DNA glycosylase | chromosome stability | 99 | *Ab, Kp, Ec, Sf, Sm* | *Po, Pm, Pa* | 1963 | no |
| PA5441 | Type IV pili protein | cell motility | 93 | *Ab, Kp, Ec, Sm, Sp, Sa* | *Po* | 2532 | no |
| PA1009 | phosphoserine phosphatase | pyrimidine salvage pathway, genome | 99 | *Ab, Kp, Sp, Ec, Ef* | *Pm, Pa* | 2535 | no |
| PA3756 | L,D-transpeptidases | peptidoglycan biosynthesis | 100 | *Ab, Kp, Ec, Sp* | *Po, Pm* | 2516 | no |
| PA1095 | flagellar chaperon | cell motility | 94 | *Ab, Kp, Ec,* | *Po, Pm* | 1107 | no |
| PA2984 | phosphorylcholine esterase | cell adhesion | 100 | *Ab, Kp, Sp, Ec* | *Po, Pm* | 1659 | no |
| PA2151 | glycogen-degrading | host-pathogen interactions | 100 | *Ab, Kp, Ef, Sp, Sm, Sa* | *Po, Pm* | 2501 | no |
| PA1640 | patatin-like phospholipase A | secreted toxin | 100 | *Ab, Sp, Ec, Kp, Sf* | *Pm, Pa* | 653 | no |

<sup>£</sup> Number of orthologs (>70% sequence identity, >95% alignment coverage) identified in strains isolated from humans with the known disease as indicated in PGD.
<sup>€</sup> Orthologs (>70% sequence identity, >95% alignment coverage) from *P. mendocina* (*Pm*), *P. otitidis* (*Po*) and *P. alcaligenes* (*Pa*) which are considered as important pathogens from *Pseudomonas* genus.
<sup>§</sup> Homologs (>35% sequence identity, >75% alignment coverage) from 13 WHO priority pathogens listed in figure 1. Gram-positive strains are indicated bold.

| Gene | Putative biochemical function | Putative virulence function | Phyre 2 confidence [%] | Conservation |  |  |  |
| --- | --- | --- | --- | --- | --- | --- | --- |
|  |  |  |  | WHO priority pathogens<sup>§</sup> | pathogenic *Pseudomonas*<sup>ε</sup> | clinical *P. aeruginosa*<sup>ε</sup> | humans |
| PA5033 | lectin | cell adhesion | 88 | Ab, Kp | Po, Pm | 2519 | no |
| PA1981 | DNA gyrase binding-protein | antibiotic resistance | 99 | Ab, **Sp**, Ec, Sm, **Sa**, Kp, St | Pm | 261 | no |
| PA3304 | efflux pump adapter protein | antibiotic resistance | 99 | Ab, Kp, Ec, Sf, Sm | Po, Pm, Pa | 2523 | no |
| PA4679 | uracil-DNA glycosylase | chromosome stability | 99 | Ab, Kp, Ec, Sf, Sm | Po, Pm, Pa | 1963 | no |
| PA5441 | Type IV pili protein | cell motility | 93 | Ab, Kp, Ec, Sm, **Sp**, **Sa** | Po | 2532 | no |
| PA1009 | phosphoserine phosphatase | pyrimidine salvage pathway, genome integrity | 99 | Ab, Kp, **Sp**, Ec, **Ef** | Pm, Pa | 2535 | no |
| PA3756 | L, D-transpeptidases | peptidoglycan biosynthesis | 100 | Ab, Kp, Ec, **Sp** | Po, Pm | 2516 | no |
| PA1095 | flagellar chaperon | cell motility | 94 | Ab, Kp, Ec | Po, Pm | 1107 | no |
| PA2984 | phosphorylcholine esterase | cell adhesion | 100 | Ab, Kp, **Sp**, Ec | Po, Pm | 1659 | no |
| PA2151 | glycogen-degrading protein | host-pathogen interactions | 100 | Ab, Kp, **Ef**, **Sp**, Sm, **Sa** | Po, Pm | 2501 | no |
| PA1640 | patatin-like phospholipase A | secreted toxin | 100 | Ab, **Sp**, Ec, Kp, Sf | Pm, Pa | 653 | no |

<sup>ε</sup> Number of orthologs (>70% sequence identity, >95% alignment coverage) identified in strains isolated from humans with the known disease as indicated in PGD.
€ Orthologs (>70% sequence identity, >95% alignment coverage) from *P. mendocina* (*Pm*), *P. otitidis* (*Po*) and *P. alcaligenes* (*Pa*) which are considered as important pathogens from *Pseudomonas* genus.
\$ Homologs (>35% sequence identity, >75% alignment coverage) from 13 WHO priority pathogens listed in figure 1. Gram-positive strains are indicated bold.

## Discussion

Comparative modelling of the target protein with sequence homolog of known structure is a useful tool for predicting the structure of hypothetical proteins. [60–62] Rarely, homology modelling is used for proteome-wide structural and functional annotation of hypothetical proteins. [63, 64] Here, we have demonstrated the applicability of multi-template homology modelling for global assignment of biochemical functions to hundreds of hypothetical proteins of human pathogen *P. aeruginosa*, to aid the identification of potential antibiotic targets. For that, we have used the Phyre2 server, which is among the most used tools for the prediction of protein structure. [21] Phyre2 applies an advanced method for detection of remote sequence homologs with high specificity and sensitivity, to build an accurate model (rmsd to the native structure of 2 - 4 Å) even for proteins with less than 20% sequence identity. [21] In contrast to the functional prediction based on pair-wise sequence alignment, which appears to be reliable only when sequence similarity is very high because duplicates of genes can diverge to have different function [65, 66], structure-based functional predictions take advantage of the fact that the protein structure stays conserved long after sequences are substantially changed [67]. Conclusively, Phyre2 functional annotation shows an advantage over the sequence-alone homology searches.

Our set of >21000 homology models for >2100 target *P. aeruginosa* PUFs generated using homologs with known structure and in most cases experimentally determined biochemical function, was refined to identify true homologs. The biochemical functions of true template-homologs were, in the automatic procedure, assigned as putative functions to hundreds of target proteins. Results of this functional prediction are available as PaPUF database (www.iet.uni-duesseldorf.de/arbeitsgruppen/bacterial-enzymology). It is likely that for some PaPUFs homology-modelling based functional assignment is misleading because it is known that during evolution from one protein may diverge several homologs with different functions. [68] Therefore, to reinforce the prediction of biochemical functions, we recommend verifying the biochemical functions by superimposing the homology model with the template-homologs and inspecting active sites or ligand binding sites. However, this remains tedious and time-consuming and therefore is not applicable for high-throughput analysis. Our approach revealed 360 PUFs with two or three putative biochemical functions, which were associated with different template structures. This is not surprising if we consider that many enzymes catalyse more than one reaction [69–71]; that many enzymes and transporter domains have annotated ligand-binding domains [72–75] and that often several distinct domains are arranged into one functional protein [76, 77].

Although the biochemical function is an important prerequisite for R&D of drugs, alone, without biological function, it is not sufficient for identifying the potential drug target. [78, 79] Therefore, the putative biological functions of the template-homologs were acquired to predict the role of PUFs for virulence and antibiotic resistance. During the selection of putative drug targets, we have considered the information from the large-scale screens [30, 32-36] about the relationship of target *P. aeruginosa* PUFs with virulence or essential-processes, conservation of PUFs in other high-priority relevant pathogens and absence of PUF homologs in humans. This led to classify 107 PUFs (63 essential, 21 virulence factor, 28 pathogen-specific) into the highest priority group for the development of novel drugs against *P. aeruginosa*. Noteworthy, 103 of these PUFs do not have human homolog, which is a good starting point for the development of selective drugs. For 41 of these top priority drug targets were assigned putative biochemical functions with high homology modelling confidence, coverage of query protein and stereochemical parameters. These functions allowed us, based on the biological processes ascribed to homology-templates, to link them with maintaining genome integrity (putative phosphoserine phosphatase PA1009 and putative uracil-DNA glycosylase PA4679), peptidoglycan biosynthesis (putative L,D-transpeptidase PA3756), host-pathogen interactions (putative extracellular lectin PA5033, putative phosphorylcholine esterase PA2984, putative flagellar chaperon PA1095, putative Type IV pili biogenesis protein PA5441 and putative glycogen-degrading protein PA2151), extracellular toxin (patatin-like phospholipase PA1640) and antibiotic resistances (putative DNA gyrase binding protein PA1981 and putative efflux pump adapter protein PA3304) (Table 1). Interestingly, several of these drug targets were previously identified as virulence-related genes using a *Rattus norvegicus* virulence assay [54] or motility assay [36], although their functions were unknown.

All these processes are common drug targets; therefore, we believe that these proteins are promising targets for further experimental characterisation and design of inhibitors, which might represent novel antibiotics to treat not only *P. aeruginosa* but also other high-priority pathogens in which these proteins are largely conserved.

## Acknowledgement

Both authors thank the Jürgen Manchot Stiftung for funding and Prof. Karl-Erich Jaeger from the Institute of Molecular Enzyme Technology (Heinrich Heine University Düsseldorf) for valuable discussions.

