## Supplemental information for "Predicting drug targets by homology modelling of *Pseudomonas aeruginosa* proteins of unknown function"

Content: three figures and eight tables

**B)**

|  |  |
| --- | --- |
| <10% | 13 |
| 10-20% | 76 |
| 20-30% | 107 |
| 30-40% | 97 |
| 40-50% | 90 |
| 50-60% | 71 |
| 60-70% | 62 |
| 70-80% | 56 |
| 80-90% | 58 |
| 90-100% | 1474 |

|  |  |
| --- | --- |
| <10% | 984 |
| 10-20% | 387 |
| 20-30% | 285 |
| 30-40% | 183 |
| 40-50% | 111 |
| 50-60% | 55 |
| 60-70% | 23 |
| 70-80% | 11 |
| 80-90% | 7 |
| 90-100% | 56 |

**Figure S1**: Confidence score **(A)** and sequence identity **(B)** distributions for homology modelling of all 2102 *P. aeruginosa* PUFs obtained with the use of Phyre2 server**.** The only highest confidence score of each PUF was considered in the analysis.

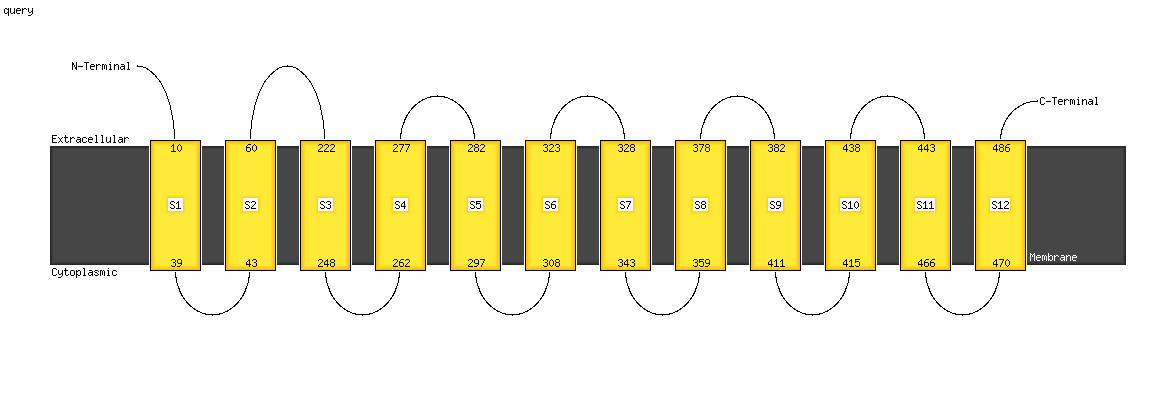

**Figure S2:** Topology of putative transmembrane helices predicted in PA2984 by Phyre2 web server.

**A)**

```
PA4679 LLVELPTG----EPFASRDPGYLLLRDLLRAA-GLPDSPRLIGEPVR
1UI0   IVGEGPGEeedkTGRPFVGKAGQLLNRILEAA-GIPREEVYITNIVK
4ZBZ   FVGEAPGEnedkEGRPFVGAAGKLLTQMIKEIlGLERDQVFITNVVK
6IOD   MIGEQPGDkedlAGLPFVGPAGRLLDRALEAA-DIDRDALYVTNAVK
4ZBX   FVGEAPGEnedkEGRPFVGAAGKLLTQMIKEIlGLERDQVFITNVVK
1VK2   FVGEGPGEeedkTGRPFVGRAGXLLTELLRES-GIRREDVYICNVVK
```

**B)**

```
PA3756 TNGCIAMNNTDMREVWSVVKDGTLIE
3VYN   SHGCLNVSPSNAQWFYDHVKRGDIVE
4XVO   SHGCINLSPDNAAWYYDXVSVGDPII
4K73   SHGCISLSREDAEWYYNAVDIGDPVI
```

**C)**

```
PA2984 LDGLLLSHADNDHAG  ~  LTGDISRAAEHAWLARQADPRVDWLLAPHHG
1WRA   LDFILVTHTHSDHIG  ~  LGGDLDNVHGAEDKYGPLIGKVDLMKFNHHH
6KNS   LDAVILSHYHHDHIA  ~  YTADSSY----QDSFIPFSENADLLISE-CN
6JKW   LTKIFLTHLHTDHWG  ~  FGGDTAP----NIWYPEYAKGADLAIH--EC
```

**Figure S3**: Conservation of functional residues among PaPUFs and their structural homologs obtained by DALI database. [1] Literature reported active site and ligand binding residues are highlighted gray. Superimposition parameters are provided in table S8.

**Table S1: Classification of *P. aeruginosa* GUFs.**

**Excel file “Table S1”**

^a^ Pathogen associated genes were obtained from PseudoCAP curation [2].

^b^ Virulence factors were obtained from Virulence factor database [3], Victors virulence factors knowledge base [4] and PseudoCAP curation [2].

^c^ General essential genes were obtained from Lee et al. (2011). [5]

**Table S2:** Priority pathogens analyzed in this study.

| Organism | Taxonomic identifier | Nr. of homologs of 41 PUFs* |
| --- | --- | --- |
| *Acinetobacter baumanii* | 470 | 41 |
| *Klebsiella pneumoniae* | 573 | 40 |
| *Escherichia coli* | 562 | 41 |
| *Serratia marcescens* | 615 | 33 |
| *Campylobacter sp.* AG18-0001 | 197 | 32 |
| *Salmonella typhi* | 90370 | 30 |
| *Staphylococcus aureus* | 1280 | 25 |
| *Neisseria gonorrhoeae* | 485 | 25 |
| *Enterococcus faecium* | 1352 | 25 |
| *Helicobacter pylori* | 210 | 16 |
| *Streptococcus pneumoniae* | 1313 | 38 |
| *Shigella flexneri* | 623 | 31 |
| *Haemophilus influenzae* | 727 | 23 |

*Virulence-related and essential PUFs with predicted biochemical functions were used as a query for BLAST search on NCBI.

**Table S3:** *P. aeruginosa* GUFs with experimentally demonstrated virulence function.

| **PGD ID** | **GUF category** | **Virulence process** | **Reference** |
| --- | --- | --- | --- |
| PA2146 | conserved hypothetical | *Hordeum vulgare* | [6] |
| PA2360 | hypothetical | *Caenorhabditis elegans* | [7] |
| PA2372 | hypothetical | *Caenorhabditis elegans* | [7] |
| PA2827 | conserved hypothetical | *Drosophila melanogaster* | [8] |
| PA1009 | hypothetical | *Rattus norvegicus* | [9] |
| PA2972 | conserved hypothetical | *Rattus norvegicus* | [9] |
| PA3756 | hypothetical | *Rattus norvegicus* | [9] |
| PA3826 | hypothetical | *Rattus norvegicus* | [9] |
| PA4115 | conserved hypothetical | *Rattus norvegicus* | [9] |
| PA4564 | conserved hypothetical | *Rattus norvegicus* | [9] |
| PA4692 | conserved hypothetical | *Rattus norvegicus* | [9] |
| PA5441 | hypothetical | *Rattus norvegicus* | [9] |
| PA2146 | conserved hypothetical | *Hordeum vulgare* | [6] |
| PA2462 | hypothetical | Hemolysis | [6] |
| PA1093 | hypothetical | Flagellum function | [10] |
| PA1095 | hypothetical | Flagelum function | [10] |
| PA1096 | hypothetical | Flagelum function | [10] |
| PA1442 | conserved hypothetical | Flagelum function | [10] |

**Table S4:** Homology modelling of *P. aeruginosa* PUFs using Phyre2 server.

Excel file “Table S4”

**Table S5:** PUFs with more than 96% of sequence identity to templates with known function.

| **PA Nr** | **Confidence (%)** | **Sequence identity (%)** | **Query coverage*** | **PDB ID of template** | **Function** | **PubMed ID** |
| --- | --- | --- | --- | --- | --- | --- |
| PA0666 | 100,0 | 100 | 360 | 3qbw | 1,6-Anhydro-N-acetylmuramic acid kinase | 21288904 |
| PA3800 | 100,0 | 100 | 352 | 4hdj | BamB, component of β-Barrel Assembly Machine (BAM) | 23189157 |
| PA3086 | 99,5 | 100 | 69 | 2gqc | protease | 17059825 |
| PA4534 | 100,0 | 100 | 136 | 4ubr | N-acetyltransferase | to be published |
| PA4279 | 100,0 | 98 | 245 | 2f9t | transferase | 16905099 |
| PA3263 | 100,0 | 100 | 305 | 2owy | recombination-associated protein RdgC | 17426134 |
| PA3302 | 100,0 | 100 | 154 | 5cpg | lyase | 26386053 |
| PA4991 | 100,0 | 98 | 390 | 5ez7 | FAD dependent oxidoreductase | 26841760 |
| PA3764 | 100,0 | 97 | 425 | 4oz9 | lyase | 27618662 |
| PA0616 | 100,0 | 100 | 174 | 4s37 | R2 pyocin membrane-piercing spike | to be published |
| PA0115 | 100,0 | 98 | 148 | 1xeb | Acyl-CoA N-acyltransferase | to be published |
| PA5396 | 100,0 | 97 | 324 | 2i5g | amidohydrolase | to be published |
| PA5201 | 100,0 | 96 | 323 | 3bzk | transcription Tex protein | 18321528 |
| PA1221 | 100,0 | 99 | 586 | 4dg9 | ligase | 22452656 |
| PA5185 | 100,0 | 100 | 138 | 2o5u | thioesterase | 19898606 |
| PA1865 | 100,0 | 100 | 537 | 4r8a | hydrolase | 25319828 |
| PA4992 | 100,0 | 100 | 267 | 4exa | Aldo_ket_red domain-containing protein | 23295481 |
| PA4872 | 100,0 | 100 | 283 | 3b8i | oxaloacetate decarboxylase | 18081320 |

*Number of residues.

**Table S6:** Keywords used for textual mining of enzyme, ligand binding and transporter functions of templates used for modelling of *P. aeruginosa* PUFs.

| **Hydrolase** | **Transferase** | **Oxidoreductase** | **Ligase** | **Lyase** | **Isomerase** |
| --- | --- | --- | --- | --- | --- |
| *hydrolase* | *transferase* | *oxidoreductase* | *ligase* | *lyase* | *isomerase* |
| *cellulase* | *kinase* | *hydroxylase* | *synthetase* | *aldolase* | *epimerase* |
| *glucosidase* | *phosphorylase* | *reductase* | *synthase* | *fumarse* | *mutase* |
| *gtpase* | *transaldolase* | *oxidase* |  | *cyclase* | *racemase* |
| *atpase* | *transglutaminase* | *oxido-reductase* |  | *dehydrochlorinase* |  |
| *peptide release* | *polymerase* | *flavoenzyme* |  |  |  |
| *peptidase* | *thiolase* | *hydrogenase* |  |  |  |
| *nuclease* |  | *oxygenase* |  |  |  |
| *dnase* |  | *cytochrom* |  |  |  |
| *rnase* |  |  |  |  |  |
| *phosphatase* |  |  |  |  |  |
| *amylase* |  |  |  |  |  |
| *amidase* |  |  |  |  |  |
| *esterase* |  |  |  |  |  |
| *protease* |  |  |  |  |  |
| *proteinase* |  |  |  |  |  |
| *helicase* |  |  |  |  |  |
| *leishmanolysin* |  |  |  |  |  |
| *hydrolytic enzyme* |  |  |  |  |  |
| *lipase* |  |  |  |  |  |
| *phospholipase* |  |  |  |  |  |
| **Binding** | | | | **Transporter** | |
| **Nucleic acids** | **Lipid** | **Sugar** | **Nucleotid** |  | |
| *dna* | *lipid* | *cellulose* | *atp* | *porin* | *export* |
| *rna* | *fatty acid* | *carbohydrat* | *amp* | *channel* | *import* |
| *nucleic* |  | *peptidoglycan* | *nucleot* | *efflux* | *uptake* |
|  |  | *lectin* | *fmn* | *pump* | *permease* |
|  |  | *sugar* |  | *transporter* | *anti/sim/porter* |
|  |  | *maltose* |  |  |  |

* indicate any character

**Table S7:** Stereochemical validation of PaPUF homology models with PROCHECK. [11]

| **PaPUF** | **Core (%)** | **Allowed (%)** | **General (%)** | **Disallowed (%)** |
| --- | --- | --- | --- | --- |
| PA2151 | 81.0 | 14.7 | 2.3 | 1.9 |
| PA2984 | 80.5 | 14.5 | 2.0 | 3.0 |
| PA5033 | 73.6 | 22.0 | 3.1 | 1.3 |
| PA1095 | 89.4 | 9.6 | 1.0 | 0.0 |
| PA3756 | 79.7 | 16.1 | 3.4 | 0.8 |
| PA1009 | 85.2 | 13.6 | 0.0 | 1.2 |
| PA4679 | 83.8 | 12.0 | 3.4 | 0.9 |
| PA3304 | 78.3 | 16.5 | 3.3 | 1.9 |
| PA1981 | 72.7 | 26.0 | 0.6 | 0.6 |
| PA5441 | 91.4 | 8.1 | 0.0 | 0.5 |
| PA1640 | 86.6 | 11.5 | 0.5 | 1.4 |

**Table S8.** Structural alignments of PUF homology model with its corresponding template and additional proteins belonging to the same functional class, according to DALI database.

| **PaPUF** | **Phyre2 template***  **PDB ID / rmsd [Å]** | **Homolog 1**  **PDB ID / rmsd [Å]** | **Homolog 2**  **PDB ID / rmsd [Å]** | **Homolog 3**  **PDB ID / rmsd [Å]** | **Homolog 4**  **PDB ID / rmsd [Å]** |
| --- | --- | --- | --- | --- | --- |
| PA2984 | 1WRA / 1.2 | 6KNS / 3.2 | 6JKW / 3.3 | - | - |
| PA3756 | 3VYN / 0.6 | 4XVO / 1.7 | 4K73 / 1.3 | - | - |
| PA4679 | 1UI0 / 0.8 | 4ZBZ / 1.4 | 6IOD / 1.7 | 4ZBX / 1.4 | 1VK2 / 1.4 |

**References to supporting information:**
